# Consistent differences among individuals more influential than energetic trade-offs in life history variation in grey seals (*Halichoerus grypus*)

**DOI:** 10.1101/2021.07.10.451927

**Authors:** Janelle J. Badger, W. Don Bowen, Cornelia E. den Heyer, Greg A. Breed

## Abstract

Life history variation is thought to be mainly a result of energetic trade-offs among fitness components; however, detecting these trade-offs in natural populations has yielded mixed results. Individual quality and environmental variation may mask expected relationships among fitness components because some higher quality individuals may be able to acquire more resources and invest more in all functions. Thus, life history variation may be more affected by variation in individual quality than varying strategies to resolve energetic trade-offs, e.g. costs of reproduction. Here, we investigated whether variation in female quality or costs of reproduction is a larger factor in shaping differences in life history trajectories by assessing the relationship between survival and individual reproductive performance using a 32-year longitudinal data set of repeated reproductive measurements from 273 individually marked, known-aged female grey seals (*Halichoerus grypus*) from the Sable Island breeding colony. We defined individual reproductive performance using two traits: reproductive frequency (a female’s probability of breeding) and provisioning performance (provisions given to young measured by offspring mass), computed using mixed effects models separately for (1) all reproductive events, and (2) an age-class specific reproductive investment. Individual differences contributed a large portion of the variance in reproductive traits, with individuals displaying a range in individual reproductive frequencies from 0.45 to 0.94, and a range of average pup weaning masses from 34.9 kg to 61.8 kg across their lifetime. We used a Cormack-Jolly-Seber open-population model to estimate the effect of these reproductive performance traits on adult survival probability. Our approach estimated a positive relationship between reproductive performance and survival, where individuals that consistently invest well in their offspring survive longer. The best supported model estimated survival as a function of age-class specific provisioning performance, where late-life performance was quite variable and had the greatest impact on survival, possibly indicating individual variation in senescence. There was no evidence to support a trade-off in reproductive performance and survival at the individual level. These results suggest that in grey seals, individual quality is a stronger driver in life history variation than varying strategies to mitigate trade-offs among fitness components.

## Introduction

Inherent trade-offs among fitness components have been shown to influence diverse biological phenomena ranging from population structure and behavior to the evolution of senescence (Hamilton 1966, Stearns 2000, Roff 2002). With limited resources to allocate, investment into a particular fitness component may result in reduced investment in others, or may come at the cost of future allocation (Cody 1966). For example, a considerable body of early theoretical work predicted the high energetic demand of reproduction should result in reduced future reproduction and/or longevity through such trade-offs (Fisher 1930, Cody 1966, Williams 1966, Partridge and Harvey 1988, Stearns 1989).

Previous research reports these trade-offs via negative associations among life history traits, e.g. lifespan and fecundity, when comparisons are made across taxonomic groups (Gaillard et al. 1989, Jones et al. 2008, Hamel et al. 2010, De Loof 2011, Lemaître et al. 2015, Salguero-Gómez et al. 2016). A wide range of species, particularly invertebrates, has evolved to maximize reproductive output over short life spans (Stearns 1992). For example, female burrowing mayflies *Hexagenia limbata* lay up to 8,000 eggs in their two-day adult life span (Hunt 1951). Though number of offspring and overall investment is large, there is minimal investment into each egg and very few survive (Hunt 1951). Longer-lived, iteroparous species may forego large quantities of offspring to invest heavily in a few offspring over longer lives (Lack 1947, Clutton-Brock et al. 1989). The longest-living mammal, the bowhead whale (*Balaena mysticetus*), is a quintessential example of this strategy (George et al. 2020). Bowhead whales do not become sexually mature until 25 years of age, and subsequently produce only one calf every 3-4 years across lifespans that can exceed ∼200 years (George et al. 1999). Contrasting life history characteristics can arise from sources such as variable predation risks (Reznick 1982) and environmental predictability (Reed et al. 2010) that influence population-level mortality patterns and resource allocation to shape natural selection (Partridge and Harvey 1988, George et al. 2020).

These optimality theories regarding trade-offs (summarized in Roff 1992 and Stearns 1992) assume homogeneity of individuals and often fail to explain life history variation among individuals when empirically tested within populations (van Noordwijk and de Jong 1986, Cam et al. 2002, Bolnick et al. 2003, De Loof 2011). Positive, zero, or negative correlations may arise between two traits that are involved in a classical physiological trade-off when measurements are analyzed on the individual scale. As an example, if individuals vary in their ability to acquire resources then we would expect those able to collect more resources to allocate more to all functions (van Noordwijk and de Jong 1986, Reznick et al. 2000), resulting in positive associations among fitness traits. Certain individuals in a population may be of higher quality, or “better”, than others, as an emergent property of endogenous (e.g. genetic differences, ontogenetic development, phenotypic traits, physiology, behavior) and exogenous (e.g. habitat, diet quality) factors or interactions among them (Reznick et al. 2000, Bolnick et al. 2003, Araújo et al. 2011, Bolnick et al. 2011). Though energetic trade-offs may be physiologically unavoidable, higher quality individuals that acquire more resources, and/or are better able to use available resources, can allocate more to both survival and reproduction and thereby appear to “override” costs of breeding (Clutton-Brock 1984, Bérubé et al. 1999, Weladji et al. 2008).

In iteroparous species, positive relationships between breeding success and survival on an individual level are well documented (Gimenez et al. 2017). However, many of these investigations involve only the trade-off between current reproduction and survival to the following year (e.g. Beauplet et al. 2006, Weladji et al. 2008, Cox and Calsbeek 2010, Blomberg et al. 2013, Stoelting et al. 2015) and are likely to underestimate the total costs of breeding (Clutton-Brock 1984). The relationship between investment and survival may be more complicated, especially for long-lived species that must carefully optimize allocation of energy to maximize fitness over many reproductive attempts (Hamilton 1966, Bell 1980, Begon and Parker 1986, Partridge and Harvey 1988, Festa-Bianchet and Jorgenson 1998). The costs associated with multiparity may accumulate over time, resulting in accelerated senescence rather than reduced survival directly following a year of high allocation of resources to reproduction (Williams 1966, Blomquist 2009). Nevertheless, trade-offs may still govern how individuals allocate resources to different essential functions over time (Clutton-Brock 1984), and an individual’s investment strategy may skew the relationship between reproductive investment and longevity. For example, heavy energy allocation into reproduction early in life may stunt growth and limit longevity independent of overall reproductive investment (Stearns 1989, Reed et al. 2008, Aubry et al. 2009, Blomquist 2009). Physiological decline associated with senescence (Williams 1957, Kirkwood and Austad 2000) may make increased reproductive investment late in life unprofitable (Hamilton 1966, Kirkwood 1977). However, regardless of total reproductive performance, this physiological decline may be delayed if early-life reproduction is also slower (Charmantier et al. 2006). In order to determine if observed positive relationships are an indication of differences in individual quality or an artifact of insufficient data to elucidate trade-offs, analysis of long time series of individual investment is needed to demonstrate costs of reproduction in long-lived animals.

The extensively studied colony of grey seals (*Halichoerus grypus*) breeding on Sable Island, Nova Scotia provide an excellent opportunity to investigate life history variation. Grey seals are long-lived (∼ 40 years), iteroparous capital breeders in which females invest heavily into the survival of a single offspring over the course of a relatively short lactation period lasting 16-18 days (Boness and James 1979, Iverson et al. 1993). During the nursing period, mothers lose a third of their body mass on average (4.1 kg per day, Mellish et al. 1999) relying only on fat reserves to produce milk and maintain metabolism, while their pups typically more than triple their birth mass (2.8 kg per day, Bowen et al. 1992). This energy transfer has a direct impact on female fitness – pups that have a higher mass at weaning are more likely to survive to recruit to the breeding population (Bowen et al. 2015). At the end of lactation, females abruptly end care, mate, and return to the sea, which conveniently allows female reproductive expenditure to be accurately measured by the energy allocated to offspring (Bowen et al. 2007). Here, we use a longitudinal data set of repeated reproductive measurements from individually marked, known-aged female grey seals to investigate whether variation in female quality or costs of reproduction are more influential in differences among individual life history trajectories.

Recent analyses indicate that typical demographic stratification such as age and sex do not adequately explain the distribution of survival rates and reproductive capabilities in a wide range of long-lived species (Bolnick et al. 2003, Cam et al. 2002, Chambert et al. 2013, Gimenez et al. 2017). This is also the case for the vital rates of adult grey seals (Bowen et al. 2007, den Heyer and Bowen 2017, Badger et al. 2020). Individuals vary substantially in reproductive success, where some females consistently outperform others on multiple metrics (Badger et al. 2020). Females that have larger pups on average tend to reproduce more often throughout their lives, and individuals that breed in a given year are more likely to breed in future years. Though female survival is high and relatively constant (0.989±0.001 for females aged 4-24, 0.901±0.004 for females aged 25+, den Heyer and Bowen 2017), life history theory predicts individual reproductive history and allocation of energy would explain some of the variation in mortality. If variation in individual quality drives differences in life history trajectories in this population, then individuals with greater reproductive success would also be those that survive longer (e.g. such as in the good genes hypothesis, Curio 1983). Alternatively, if varying strategies to mitigate reproductive costs shapes variation in life histories, individuals may invest less in each reproductive event to live longer and produce a greater number of offspring. Thus our long-term data set allows us to test the relative strength of the two most prominent sources predicting inter-individual variation in reproductive performance: individual quality (Cam et al. 2002, Bolnick et al. 2003, Bergeron et al. 2011), and reproductive trade-off, which is now best articulated in the pace-of-life syndrome hypothesis (POLS; Roff 2002, Jones et al. 2008)

To test these hypotheses for drivers of life history variation in the Sable Island breeding colony, we analyzed 32 years of mark-recapture data and reproductive allocation data to assess the relationship between survival and individual reproductive performance. We explored this relationship for both lifetime reproductive investment and age-class-dependent investment strategies. If high individual performance is associated with decreased survival (negative relationship), we consider that support for a *costs of reproduction* hypothesis (consistent with POLS), while a positive relationship between individual performance and survival would indicate support for an *individual quality* hypothesis.

## Methods

This study was conducted on Sable Island, Canada (43.93°N, 59.91°W), a partially vegetated sandbar on the Scotian Shelf roughly 160 km off the coast of Nova Scotia, Canada, during the 1989-2020 breeding seasons. The breeding season at this colony spans early December through early February, with 97% of pups born by mid-January (Bowen et al. 2007, den Heyer et al. 2017). Sable Island supports the largest breeding colony of grey seals in the world with an estimated 83,600 pups born on the island in 2016 (den Heyer et al. 2017).

### Data Collection

We used a 32-year mark-resighting dataset (1989-2020) of known-aged grey seal females to determine how lifetime reproductive performance influences survival. Individuals were marked as pups shortly after weaning with unique alpha-numeric hot-iron brands in 1985, 1986, 1987, and 1989. These permanent brands allowed reliable identification of individuals over the course of their lives. Females can recruit to the breeding population as early as 4 years old, but this is uncommon, and the average age of first reproduction is 5.6 ± 0.12 SE years for these cohorts (den Heyer et al. 2013). Each breeding season since 1989, teams of researchers conducted 5-7 roughly weekly censuses of branded females returning to the island to give birth and mate. Once sighted, branded individuals with dependent pups were visited daily from a roughly 30m distance. Prior to weaning, pups were sexed and marked with semipermanent, uniquely numbered tags in the hind flipper to ensure accurate identification after the marked female ended lactation and returned to sea, leaving her pup in the colony. Females attend their pups continuously throughout lactation. Therefore, once a pup was sighted alone, it was considered weaned and weighed to the nearest 0.5-kg.

The probability of observing a marked female during any given year includes both the probability the female is present, and the probability that she is detected given presence at the breeding colony. Females not seen in any of the weekly brand-resighting censuses in a given year could arise via one of three events: 1) the female died, 2) the female was not present at the breeding colony due to temporary emigration (“reproductive skipping”) or they quickly abandoned their pup, or 3) was present in the breeding colony, gave birth and lactated normally, but was not detected during weekly censuses either because of her position on the island or unreadable brand markings. For individuals branded from 1998-2002, 3.7% or 170 of 4569 sightings (from breeding seasons 2002 to 2012) were not readable either because of a poor-quality brand or poor conditions (Bowen et al. 2015). Low sighting probability associated with poor brands would obscure female quality, so we removed individuals whose brands were sighted as unreadable for more than 25% of their total sightings. Of the remaining brands, a recent CMR analysis of this population indicated that, if a female rears a pup on the island, there is less than a 5% chance researchers will fail to detect her in at least one resighting census (Badger et al. 2020). Grey seals are highly site philopatric, and once recruited to a breeding colony, will very rarely pup elsewhere (Bowen et al. 2015). Thus, we are able to reliably follow the reproductive history of individuals, and do not expect permanent emigration to other colonies to be a significant source of sighting error. We removed sightings of females without pups from this analysis. Individuals that are not rearing pups can be skittish and may flee to the water, resulting in a lower sighting probability than females nursing and defending young. Pregnant females generally arrive at the breeding colony five days before parturition and can travel large distances before settling to give birth (median: 2.4 km, Weitzman et al. 2017). The distance moved by nursing females from one day to the next averages about 5 m (Boness and James 1979, Weitzman et al. 2017), so movement by females during lactation is not a source of sighting error. Nursing typically lasts 2-3 weeks, so females are usually available to be resighted during two or more of the weekly re-sighting censuses in a given year.

Individual sighting histories were collected from age at first reproduction (first sighting in breeding colony) until death (or 2020 for animals still living). Sighting histories of individuals were scored as a 0 (not sighted) or 1 (sighted) for each year from 1989 to 2020. Females sighted in only one breeding season were omitted from this analysis to ensure that they had in fact recruited to the Sable Island breeding population and we have adequate data to estimate lifetime reproductive performance.

### Statistical Analysis

In this analysis, individual sighting histories and associated repeated measures of pup weaning mass were used to determine how reproductive energy allocation influences survival. Reproductive energy allocation here is defined by two traits, detailed below: (1) the energy delivered to young integrated over reproductive events, termed provisioning performance (*PP*), and (2) how often a female breeds, termed reproductive frequency (*RF*). After controlling for known effects on these variables, we estimated both reproductive traits over all reproductive events, denoted *life* and over three age-classes operationally defined as early (recruitment to age 14), peak (aged 15-24) and late (aged 25+) per previous work on age-related reproductive ecology of this species (Bowen et al. 2006, Lang et al. 2011, Bowen et al. 2015). We modeled survival as a function of reproductive investment metrics with a Cormack-Jolly-Seber open-population model. Provisioning performance, reproductive frequency, and survival were modeled simultaneously. This overall modeling framework is conceptually outlined in Figure 1, and all parameter and variable definitions are listed in Table 1 for reference.

**Table 1:**
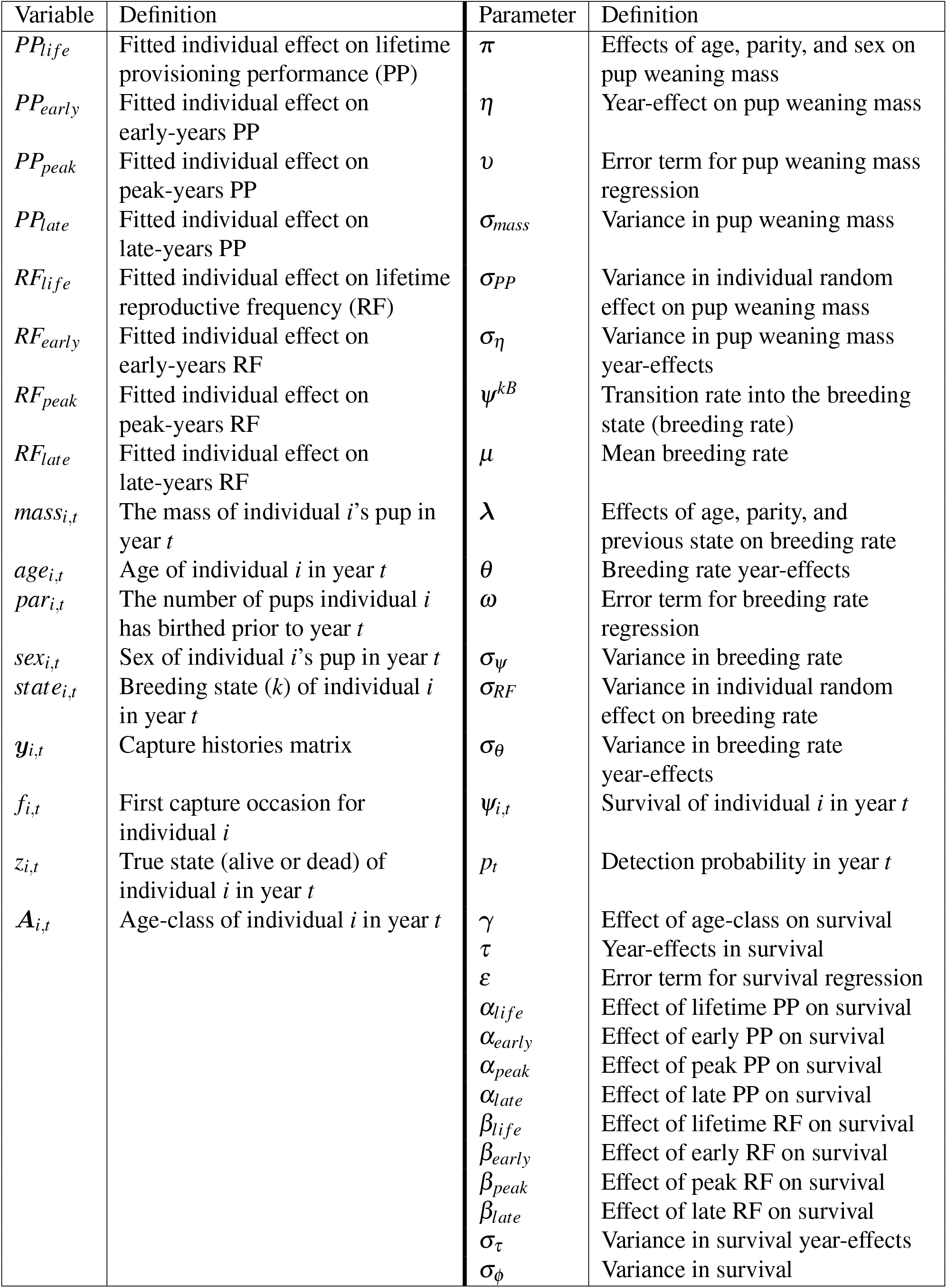
Parameter and variable definitions.

**Table 1:** WAIC estimates for 5 competing models. Model 0 is our null model, with no effect of reproductive performance on survival. Model 1 describes survival as a function of lifetime provisioning performance, Model 2 instead uses the age class effect of provisioning performance, Model 3 models survival as a function of lifetime reproductive frequency, and Model 4 describes the effect of reproductive frequency on survival as age class-specific. Estimated parameter posterior distributions from best supported model (Model 2) are found in Table 2, and distribution estimates from other models found in Appendix 1: Table S1.

| Model | Effect of Reproductive Performance | WAIC | $\Delta$ WAIC |
| --- | --- | --- | --- |
| Model 2 | Age-class provisioning performance | 2446.8 | 0 |
| Model 1 | Lifetime provisioning performance | 2449.0 | 2.2 |
| Model 4 | Age-class reproductive frequency | 2457.7 | 10.9 |
| Model 3 | Lifetime reproductive frequency | 2474.1 | 27.3 |
| Model 0 | None | 2495.6 | 48.8 |

**Figure 1:**
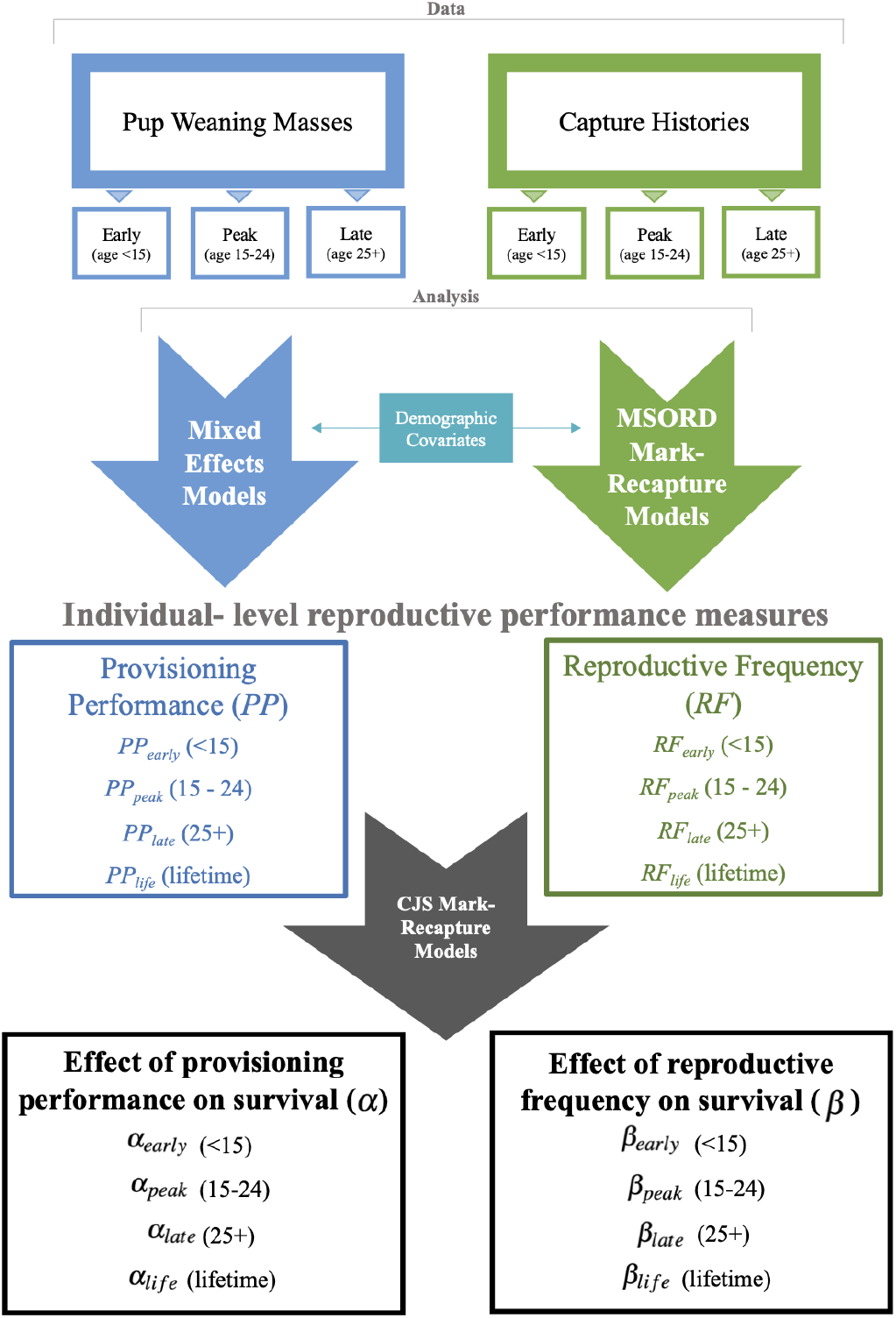
Modeling framework. Repeated measurements of pup weaning masses are discretized into full histories (*life*) and early (<15), peak (15-24) and late (25+) years then fit separately to linear mixed effects models with a quadratic age effect, pup sex, and female experience as fixed effects and individual and year as random intercepts, yielding *PP*_*life*_, *PP*_*early*_, *PP*_*peak*_, *PP*_*late*_ as predicted estimates of individual intercepts from all four models. Similarly, individual sighting histories are discretized into age-class and full histories, then separately fit to a multistate open robust design mark-recapture model (MSORD) to estimate transitions into a breeding state, with a quadratic age effect, previous state, and female experience as fixed effects, and individual and year as random effects, yielding *RF*_*life*_, *RF*_*early*_, *RF*_*peak*_, *RF*_*late*_ as predicted estimates of individual intercepts effects from all four models. These individual effects are then used as covariates in a Cormack-Jolly-Seber open population model to estimate the effect of individual reproductive performance on survival, with parameter estimates ***α*** = {*α*_*life*_, *α*_*early*_, *α*_*peak*_, *α*_*late*_} for lifetime, early, peak, and late effects of provisioning performance, respectively, and ***β*** = {*β*_*life*_, *β*_*early*_, *β*_*peak*_, *β*_*late*_} for lifetime, early, peak, and late effects of reproductive frequency, respectively.

#### Defining lifetime and age class-specific provisioning performance (PP)

During lactation, grey seal pups consume only milk provided by the female, and as capital breeders, females fast for the entire lactation period. Therefore, in our study, the body mass of a pup at weaning is a reasonable estimate of the energy (i.e. nutrients) transferred to young. While provisioning performance or maternal investment may be better measured by quantifying the change in offspring mass and/or the change in maternal mass from birth to weaning, such an endeavor is logistically limited for a longitudinal study of this scale. By using pup weaning mass as a proxy for maternal investment, we are assuming that the bulk of the variation in pup weaning masses stems from nursing and maternal behavior, and that the variation from other sources is comparatively negligible. *PP* was estimated as the average of the weaning masses of all her pups during the course of our study after accounting for confounding effects of age, experience, and offspring sex. We included age at first reproduction as a covariate as it is an important life history trait that may influence maternal investment strategies. We modeled the weaning mass of pup *j* born to of female *i* in year *t* (*mass*_*j,t*_) as a linear mixed-effects model with female experience (parity, i.e. *par*, because this effect tends to plateau, it was discretized into 1, 2, 3 and 4+ parities), pup sex, a quadratic effect of standardized female age (Bowen et al. 2006), and age at first reproduction as covariates along with random individual and year intercepts:

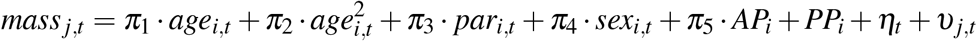

Where parameters ***π*** = {*π*_1_, *π*_2_, *π*_3_, *π*_4_, *π*_5_} reflect the quadratic age effect, effect of female experience, pup sex, and age at first reproduction, respectively, *PP*_*i*_ is the random effect of individual such that 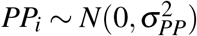, *η*_*t*_ reflects the random year effect, where 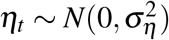, and *υ*_*j,t*_ is the error term where 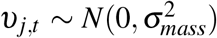. Fitted estimates of the random effect of each female (i.e. individual intercepts *PP*_*i*_) provide an estimate of that individual *i*’s performance relative to other females in the study after accounting for fixed effects. For example, a positive fitted intercept *PP*_*i*_ indicates that individual *i* on average produces larger pups at weaning than the population mean. We fit this model to the weaning masses of pups produced over the course of the female’s lives and also separately to the pups produced in the three discretized periods of the female’s lives, yielding performance metrics *PP*_*life*_, *PP*_*early*_, *PP*_*peak*_, and *PP*_*late*_, respectively. We standardized these metrics for easier interpretation of results, where the standardized metric for each of the individuals *i* = 1, …, *n* is defined as 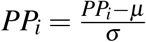 where 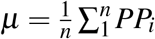 and 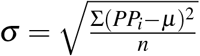. Models were validated using diagnostic plots of normalized residuals.

#### Defining lifetime and age-class specific reproductive frequency (RF)

The second reproductive trait, reproductive frequency, is defined as the probability an individual will return to the island to breed in any given year. We estimated a female’s reproductive frequency by modeling her reproductive history as a Markov chain in a multi-state open robust design capture recapture modeling framework (MSORD, following Badger et al. 2020) to account for imperfect detection in reproductive rate. Between her first and last sightings on the island during our study, a female transitions among three reproductive states: initially a first time breeder *F*, then switching between a breeder state *B*, or non-breeder state *NB*. Reproductive frequency is then defined as the probability of transition *ψ*^*kB*^ into a breeding state *B* from any reproductive state *k*. An individual’s state transition from year *t* to *t* + 1 is modeled as a categorical trial with probabilities of transition *ψ*^*ks*^ from any state *k* to any state *s, k, s* ∈ {*F, B, NB*}. We used mixed-effects logistic regression embedded in the MSORD to account for standardized female age, female experience, previous breeding state, age at first reproduction, and random individual and year effects in probability of breeding (*ψ*^*kB*^) to estimate individual intercepts:

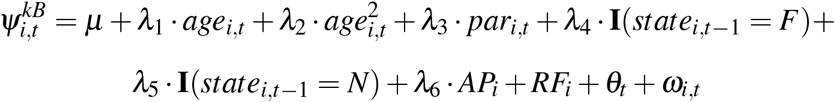

Where parameters ***λ*** = {*λ*_1_, *λ*_2_, *λ*_3_, *λ*_4_, *λ*_5_, *λ*_6_} represent the quadratic age effect, effect of female experience, the effects of the previous state, and age at first reproduction, respectively, where **I** denotes an indicator variable. *RF*_*i*_ is the random effect of individual such that 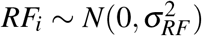, *θ*_*t*_ reflects the random year effect, where 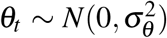, and *ω*_*i,t*_ is the error term where 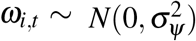. As for provisioning performance, the fitted estimate *RF*_*i*_ provides an estimate of individual *i*’s reproductive frequency relative to other females in the study. So, if a female *i*’s fitted intercept *RF*_*i*_ is negative, she reproduces less frequently than the population average after accounting for all fixed effects. We fit this model with capture histories over the course of the females’ lives and for the three age-classes separately, yielding performance metrics *RF*_*life*_, *RF*_*early*_, *RF*_*peak*_, and *RF*_*late*_, respectively. We standardized these metrics for easier interpretation of results similarly to above.

#### Survival model

We determined whether *PP* and *RF* influenced survival using a state-space formulation of the Cormack-Jolly-Seber open-population model fit to female sighting histories (Gimenez et al. 2007, Royle 2008, Kery and Schaub 2012). Our capture history matrix ***y*** has dimension *IxT*, where *I* is the total number of marked and recruited individuals and *T* is the number of years in this study. We condition on first capture (defined for individuals in vector ***f***), so recruitment is not estimated here. We define the latent state variable *z* which takes the value 1 if individual *i* is alive at time *t* (*t* = 1, …, *T*) and value 0 if the individual is dead, thus *z* defines the true state of individual *i* at time *t*. The individual is necessarily alive at its first sighting (*f*_*i*_), but states of subsequent occasions are modeled as Bernoulli trials. Conditional on being alive at occasion *t*, individual *i* may survive to occasion *t* + 1 with probability *ϕ*_*i,t*_ (*t* = 1, …, *T* -1). The Bernoulli success parameter is the product of the survival probability *ϕ*_*i,t*_ and the state of the individual *i* in time *t*, ensuring dead individuals (*z*_*i,t*_ = 0) remain dead.

However, in this and many field studies our observation of this state process is imperfect. If individual *i* is alive at occasion *t*, they may be recaptured with probability *p*_*t*_ (*t* = 2, …, *T*). So, our observations *y* are modeled as Bernoulli trials dependent on the state process *z*, where, similarly to above, the Bernoulli success parameter includes the latent state variable *z* to ensure dead individuals cannot be observed and are no longer included in the analysis. So, the state-space formulation of the CJS mark-recapture model can be defined as:

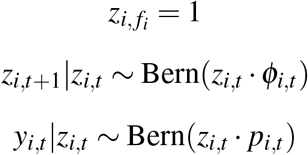

We applied a logistic regression embedded in the CJS model on survival probability *ϕ*_*i,t*_ to test our hypotheses about the effect of reproductive investment on survival. Survival probability was modeled as a function of a combination of covariates including lifetime and age class (*early, peak, late*) performance for both traits, standardized age, and a random year effect. Model 0 is our null model, with no effect of reproductive performance on survival. Model 1 includes the effect of lifetime provisioning performance *PP*_*life*_, Model 2 instead uses the age class effect of provisioning performance *PP*_*early*_, *PP*_*peak*_ and *PP*_*late*_, Model 3 includes the effect of lifetime reproductive frequency *RF*_*life*_, and Model 4 instead uses the age class effect of reproductive frequency *RF*_*early*_, *RF*_*peak*_ and *RF*_*late*_.

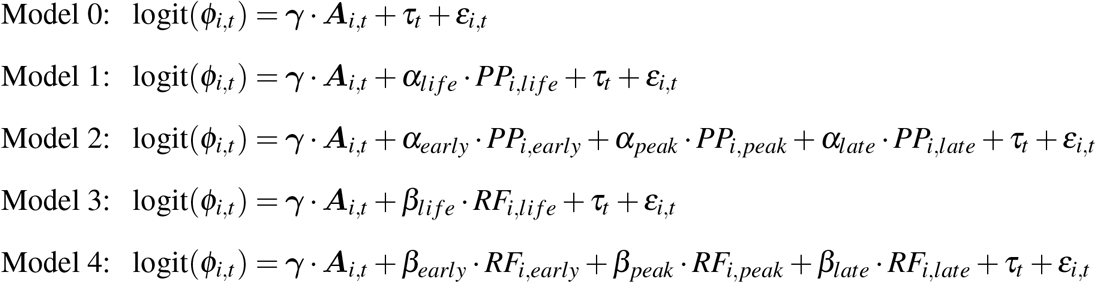

***A*** is a vector of indicator variables defining the age-class of individual *i* in time *t* (I(*age*_*i,t*_ < 15), I(*age*_*i,t*_ ≥ 15 & *age*_*i,t*_ < 25), I(*age*_*i,t*_ ≥ 25)) and ***γ*** is a vector (*γ*_1_, *γ*_2_, *γ*_3_) corresponding to mean survival rates in each age class, per previous work on grey seal survival estimates (den Heyer and Bowen 2017). *τ*_*t*_ is a random year effect such that 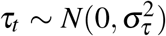, and *ε*_*i,t*_ is an error term such that 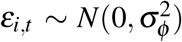. Parameters ***α*** = {*α*_*life*_, *α*_*early*_, *α*_*peak*_, *α*_*late*_} describe the effect of lifetime, early, peak, and late provisioning performance, respectively, and parameters ***β*** = {*β*_*life*_, *β*_*early*_, *β*_*peak*_, *β*_*late*_}, respectively describe the effect of lifetime, early, peak, and late age class reproductive frequency.

#### Model specification, fitting, and software

A Bayesian framework was used for model fitting, selection, and inference using the software JAGS 4.2.0 through the R interface `rjags` (Plummer 2003, R Core Team 2020, Plummer 2018). For the survival model, diffuse normal priors N(0,1000) were assigned to parameters describing the effects of age ***γ*** and components of reproductive performance ***α*** and ***β***. An uninformative Uniform(0,1) was used as a prior for probabilities of sighting *p*_*t*_. Year-effects were drawn from 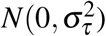 where Uniform(0,10) priors were assigned for the standard deviation in year-effects 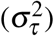. The reproductive frequency MSORD was specified following Badger et al. 2020. In the provisioning performance and reproductive frequency regression models, covariate parameters ***π*** and ***λ*** were assigned diffuse normal prior distributions *N*(0, 1000). Random year terms *η* and *θ* were specified hierarchically following a normal distribution, 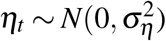 and 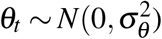, and individual terms *RF*_*i*_ were pulled from 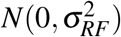. We specified a Unif(0,10) prior for random effect variance terms *σ*_*η*_, *σ*_*θ*_, *σ*_*PP*_ and *σ*_*RF*_.

Markov chain Monte Carlo (MCMC) methods were used to sample the posterior distributions of the parameters of interest. For each of the five competing models, we ran three chains in parallel using package `dclone` (Solymos 2010) with different sets of initial values. The first 10,000 MCMC samples were discarded, known as the burn-in period, after having checked that convergence was satisfactory. Convergence of chains to stationary distributions was visually evaluated using sample path plots in conjunction with the Brooks-Gelman-Rubin diagnostic 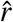 (Brooks and Gelman 1998), with values close to 1.00 indicating adequate convergence. Chains then ran for 20,000 iterations after burn-in, and a total of 2,000 MCMC samples were used for inference. We determined that a covariate had an effect if a 95% credible interval (CRI) of the posterior distribution of that parameter did not include 0. We assessed support for inclusion of reproductive performance using a measure of out-of-sample predictive ability of each model, the Widely Applicable Information Criterion (WAIC, Watanabe 2010), where a model with a smaller WAIC is judged a better fit.

## Results

We analyzed sighting histories of 273 females born in the 1985-1989 cohorts, ages 4-35, of which 16.8% (46/273) were seen at the most recent breeding season (2020), and 58.9% (161/273) and 82.1% (224/273) were seen in the last 5 and 10 years, respectively. All individuals included in this analysis were seen in all three age-classes. So, our data are right truncated with some individuals still alive and breeding. Individuals varied substantially in lifetime provisioning performance with differences among individuals explaining 45.8% of the total variation in pup weaning mass after accounting for fixed effects, resulting in a range of average pup weaning masses from 34.5 kg to 64.1 kg (individual intercepts ranging from -17.6 to 11.9 before standardizing, *PP* variance *σ*_*PP*_ = 4.87). Similarly, individuals varied in lifetime reproductive frequency, resulting in a range of lifetime reproductive frequencies from 0.45 to 0.94 (individual intercepts ranging from -1.34 to 1.28 before standardizing, *RF* variance *σ*_*RF*_ = 0.31)

Differences among individuals accounted for 45.2% of the variation in pup weaning mass in early reproduction (*PP*_*early*_ variance 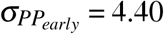), 56.6% of the variation in pup weaning mass in peak reproduction (*PP*_*peak*_ variance 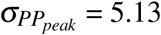), and 52.3% of the variation in late reproduction (*PP*_*late*_ variance 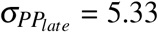). Average pup weaning masses ranged from 31.9 kg to 56.3 kg in early years, 28.4 kg to 65.1 kg in peak years, and 33.7 kg to 57.6 kg in later years. Variability among individuals in reproductive frequency was greatest in older ages (*RF*_*late*_ variance 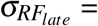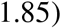, followed by peak years (*RF*_*peak*_ variance 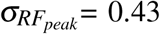) and lowest in early reproduction (*RF*_*early*_ variance 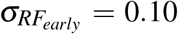). Mean reproductive frequency ranged from 0.66 to 0.79 for females in early years, 0.40 to 0.97 in peak years, and 0.11 to 0.80 later in life. Correlation among age-specific individual effects in provisioning performance were positive and quite high (Figure 2), suggesting individuals provision similarly throughout life. Correlations among individual effects in reproductive frequency were also positive, but of much lower magnitude (Figure 2).

**Figure 2:**
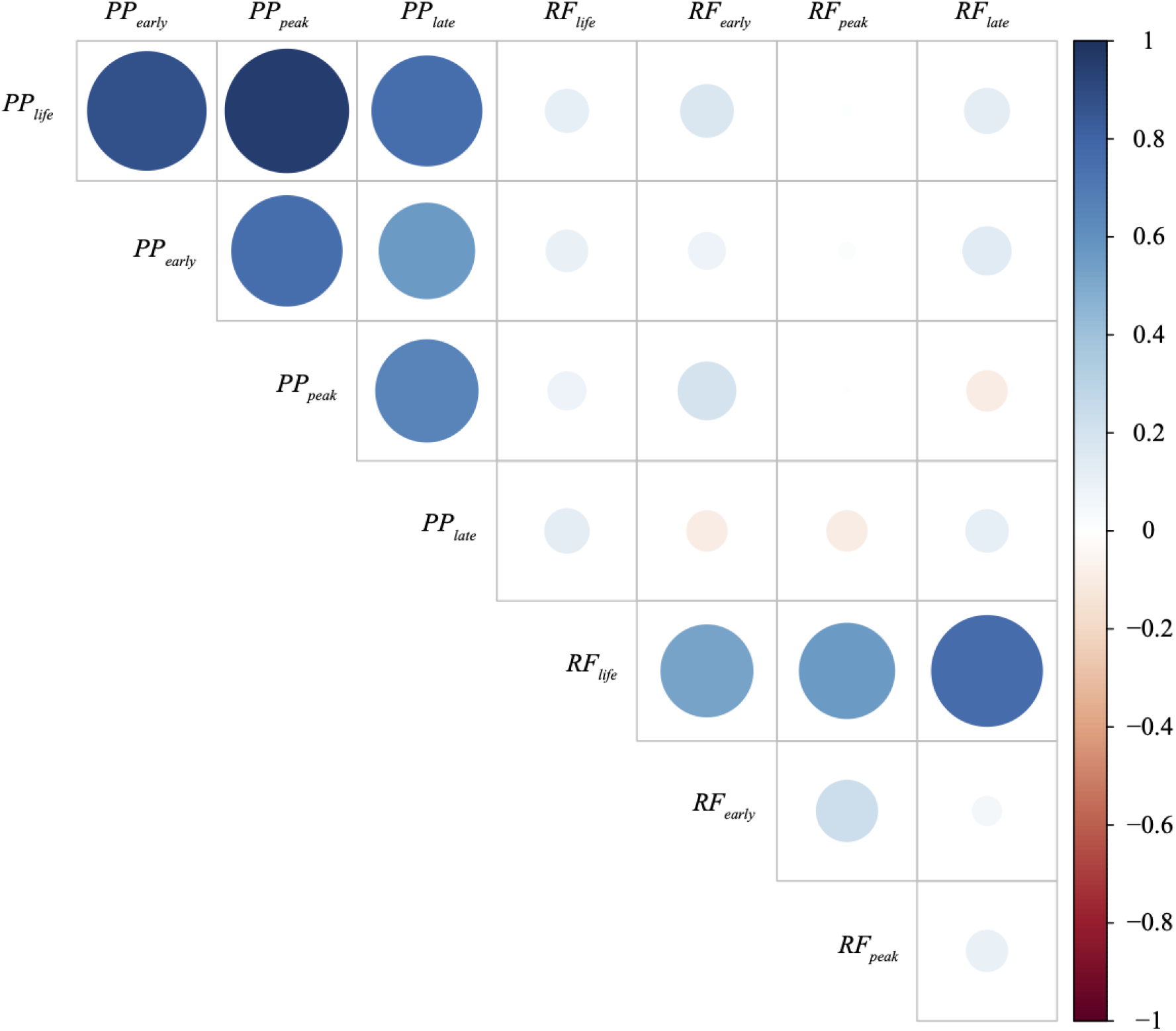
Correlation among individual effects in provisioning performance (*PP*) reproductive frequency (*RF*), where variables denoted *li f e, early, peak*, and *late* indicate lifetime, early, peak, and late performance, respectively. Blank squares indicate no correlation.

For both traits, all models agreed that individual reproductive investment has a positive effect on survival (Table 2, Appendix S1: Table S1), supporting the hypothesis that individual quality is more influential to life history variation than reproductive costs. Adult survival is particularly high and nearly invariant (den Heyer and Bowen 2017); therefore, we did not anticipate a large effect size or dramatic change in explanatory power with the inclusion of reproductive performance when modeling survival. The best supported model described survival as a function of an age class-specific provisioning performance (Table 1). While early and peak performance had negligible effects, females performing well late in life (relative to others) were more likely to survive longer. Differences in late life reproductive investment resulted in estimated survival probabilities at older ages ranging from 0.64 for the smallest *PP*_*late*_ to 0.97 for the largest *PP*_*late*_ (Figure 3). Reproductive frequency for all three age classes and lifetime performance were estimated to have positive or neutral effects on survival (Appendix S1: Table S1 & Figure 4). For both reproductive frequency and provisioning performance, late life investment had the greatest impact on survival, with higher investments associated with higher survival (Table 2, Appendix S1: Table S1).

**Table 2:**
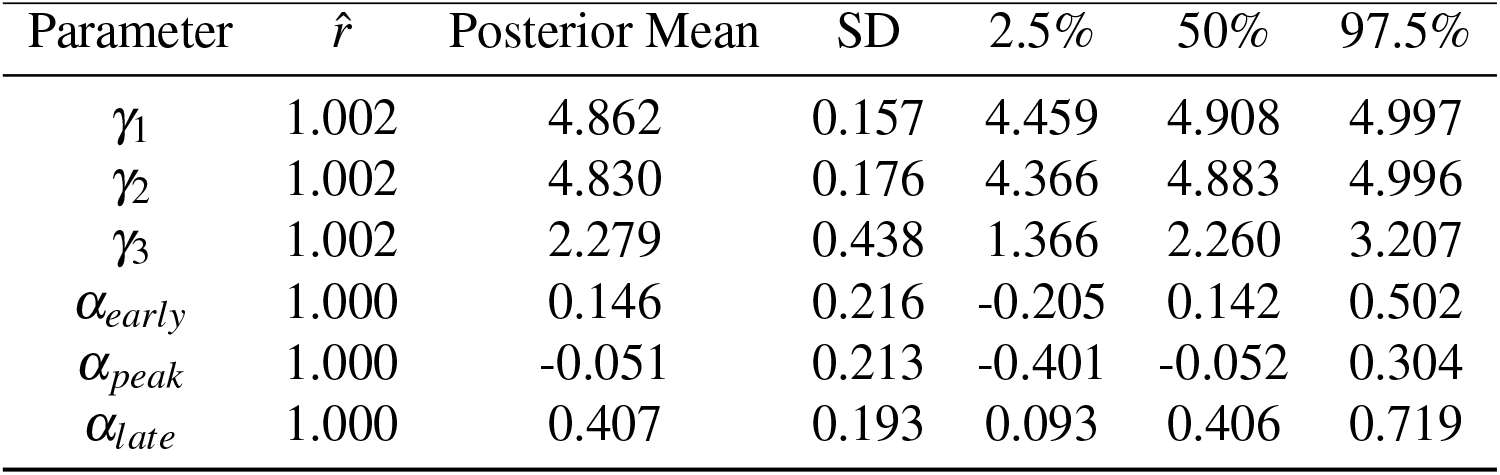
Parameter estimates from the best supported model, with survival as a function of age and individual life-age class-specific provisioning performance (Table 1). *γ*_1_, *γ*_2_, and *γ*_3_ are the age-specific survival probabilities (ages < 15, ≥15 to ≥24, and 25, respectively) and *α*_*early*_, *α*_*peak*_, and *α*_*late*_ are parameters describing the effect of early, peak, and late life provisioning performance on survival.

**Table 3:** Estimated parameter mean and 95% credible intervals in candidate models.

| Parameter | Model 0 |  | Model 1 |  | Model 3 |  | Model 4 |  |
| --- | --- | --- | --- | --- | --- | --- | --- | --- |
|  | Mean | 95% CRI | Mean | 95% CRI | Mean | 95% CRI | Mean | 95% CRI |
| $\gamma_1$ | 4.879 | [4.440, 4.978] | 4.865 | [4.497, 4.997] | 4.919 | [4.716, 4.998] | 4.924 | [4.732, 4.998] |
| $\gamma_2$ | 3.804 | [3.418, 4.238] | 4.829 | [4.363, 4.995] | 3.830 | [3.452, 4.231] | 3.948 | [3.541, 4.373] |
| $\gamma_3$ | 2.163 | [1.811, 2.601] | 2.282 | [1.470, 3.202] | 2.193 | [2.008, 2.641] | 2.208 | [2.008, 2.663] |
| $\alpha_{life}$ | - | - | 0.429 | [0.203, 0.664] | - | - | - | - |
| $\beta_{life}$ | - | - | - | - | 0.363 | [0.142, 0.582] | - | - |
| $\beta_{early}$ | - | - | - | - | - | - | 0.244 | [-0.044, 0.527] |
| $\beta_{peak}$ | - | - | - | - | - | - | 0.162 | [-0.127, 0.445] |
| $\beta_{late}$ | - | - | - | - | - | - | 0.316 | [0.097, 0.539] |

**Figure 3:**
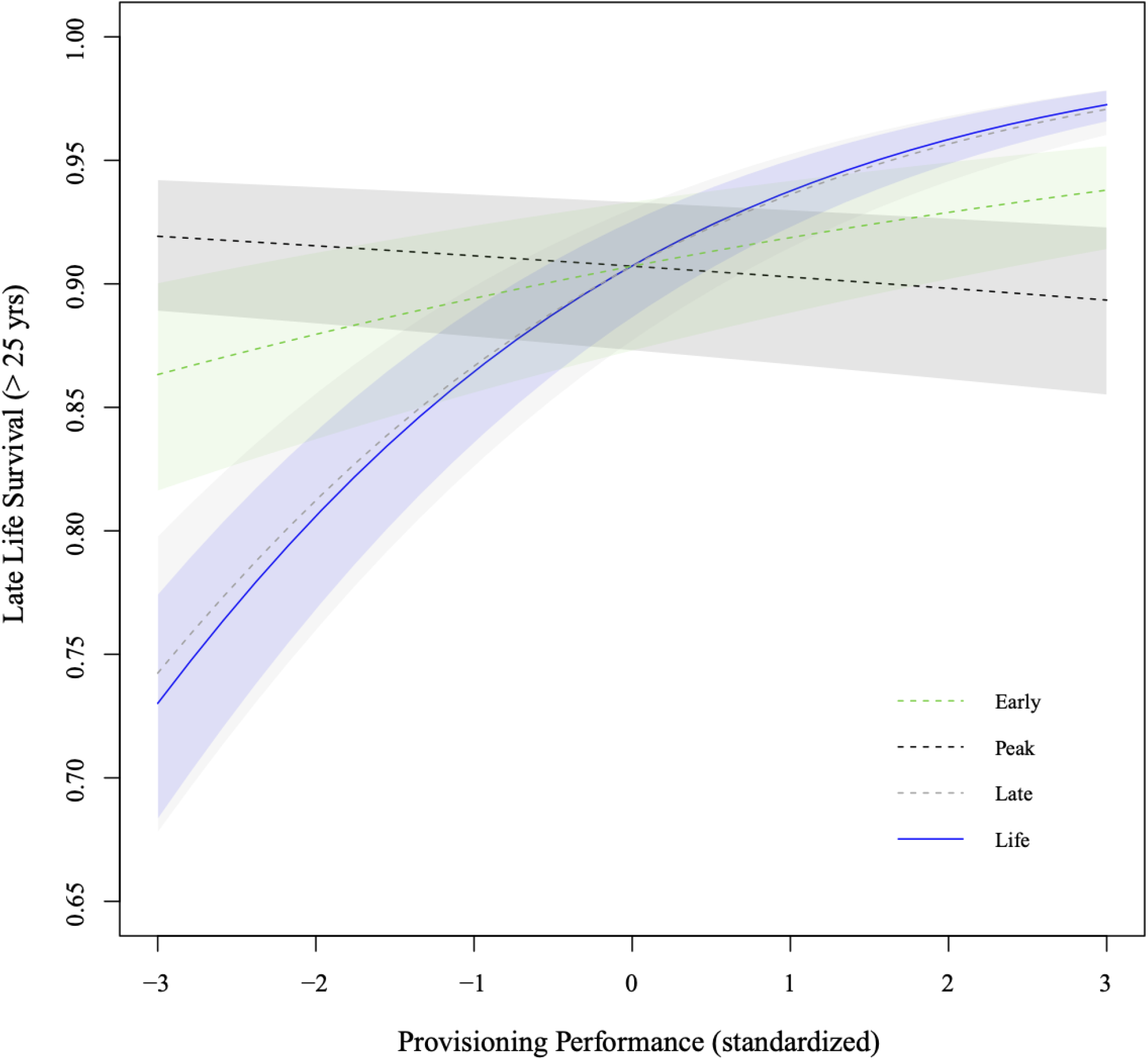
Estimated relationship between provisioning performance and late life survival (> 25 yrs old). Provisioning performance is standardized so that 0 represents the population mean. Solid blue line is relationship between survival and lifetime reproductive provisioning performance, dashed lines are age-class specific provisioning performance: green dashed line is the effect of early life investment (aged < 15 yrs), dashed black line is the effect of investment during peak years (aged 15-24 yrs), and dashed grey line is the effect of late life investment (aged 25+ yrs). Shaded regions are 95% credible intervals.

**Figure 4:**
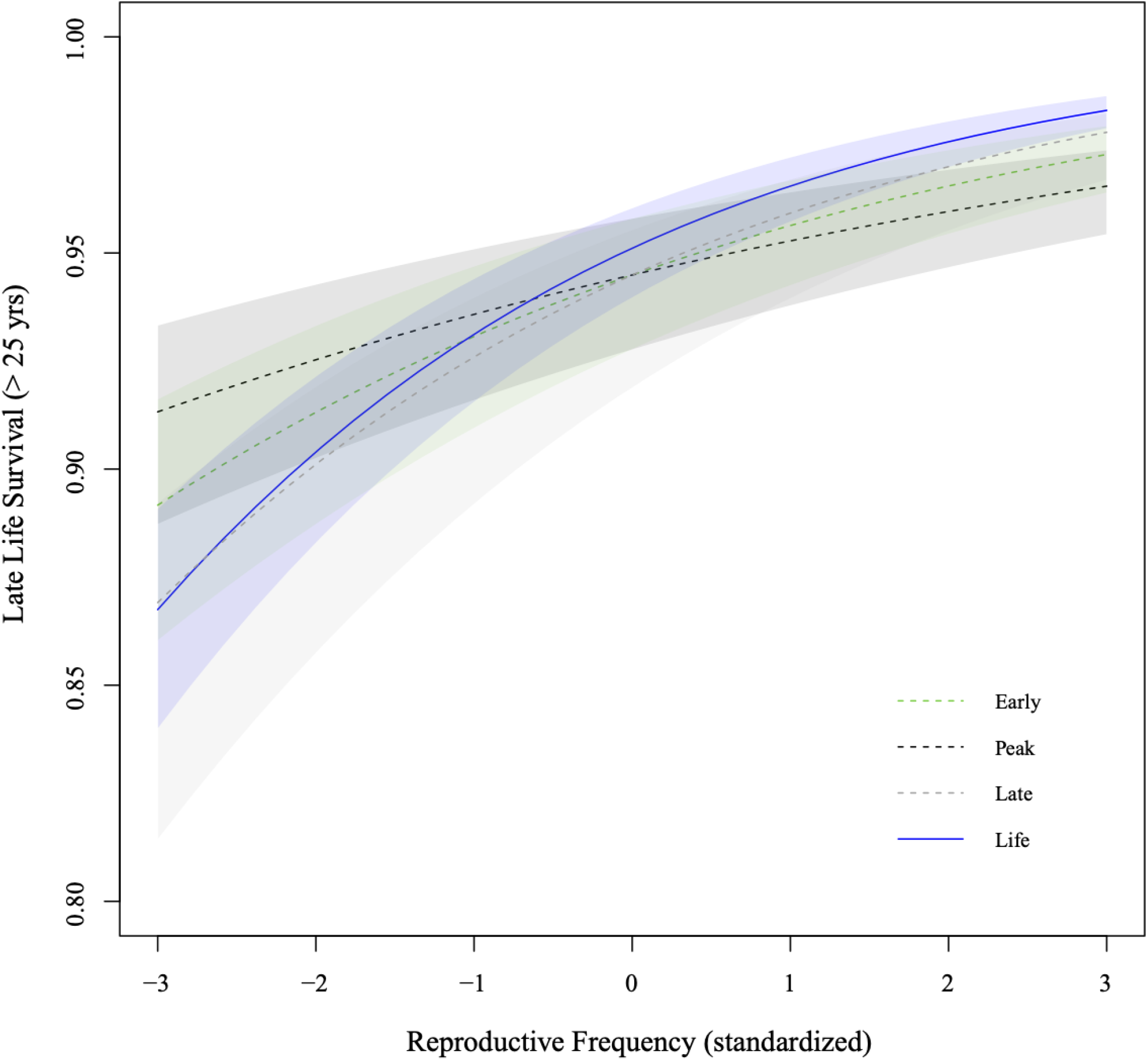
Estimated relationship between reproductive frequency and late life survival (> 25 yrs old). Reproductive frequency is standardized so that 0 represents the population mean. Solid blue line is relationship between survival and lifetime reproductive reproductive frequency, dashed lines are age-class specific reproductive frequency: green dashed line is the effect of early life investment (aged < 15 yrs), dashed black line is the effect of investment during peak years (aged 15-24 yrs), and dashed grey line is the effect of late life investment (aged 25+ yrs). Shaded regions are 95% credible intervals.

## Discussion

Here we demonstrate a clear, positive covariation between survival and both lifetime and age class-specific reproductive performance using over three decades of longitudinal data on a large sample of female grey seals. This is consistent with among-individual variation in life history traits being largely explained by differences in individual quality. Females that invested more resources to offspring and gave birth more frequently, while accounting for age, offspring sex, and environmental conditions (year effects), survived longer than females that invested less. Further, the pattern holds true across age classes, such that increased investment, particularly in later years, is associated with similar or higher survival rates. Previous work on this population has shown individual consistency in performance in multiple reproductive traits across demographic conditions (Badger et al. 2020). Taken together, these results suggest that individual quality, rather than energetic trade-offs among fitness components, is the main driver in life history variation in this population.

Positive relationships between *PP* and *RF* and subsequent survival or reproductive performance do not indicate that breeding has no reproductive costs, rather individual quality may be a more important factor in the distribution of life history phenotypes within this population which has been increasing in abundance throughout the time period of this study. Because grey seals are capital breeders that rely heavily on fat reserves, these results are likely explained by individual variation in the ability to acquire, store, and efficiently use energy sources (Bolnick et al. 2011, Araújo et al. 2011). Differences in resource acquisition may be directly linked to physical, physiological, or behavioral attributes that determine diving and foraging characteristics (Thompson and Fedak 1993, Williams et al. 2000). In this population, different individuals adopt different strategies for the timing, duration, and location of foraging bouts (Austin et al. 2004, Breed et al. 2009, Lidgard et al. 2020) which influences the foraging efficiency and effective resource availability experienced by the individuals. Positive covariances among vital rates may then derive from individuals that forage in poorer areas having both lower survival and lower reproductive success than those in better habitats.

In mammals, females also vary in their capacity to effectively transfer milk energy to offspring. Mobilization of nutrients and blubber, milk content, lactation duration, and lactation efficiency are known to vary among individuals and have important impacts on reproductive performance (Mellish et al. 1999, Lang et al. 2006, Lang et al. 2009, Crocker et al. 2014) and implications for resource allocation. Heavier mothers have more resources to allocate to milk production leading to heavier offspring at weaning (Iverson et al. 1993, Mellish et al. 1999, Macdonald et al. 2020). Maternal mass has been shown to be a dominant axis of individual quality in large mammals (Hamel et al. 2009, Plard et al. 2014), and an important determinant in yearly reproductive success in phocids (Arnbom et al. 1997, Mellish et al. 1999, Macdonald et al. 2020). Inclusion of female mass as a covariate in these analyses could lend evidence as to whether maternal energy stores help predict the variation in life histories; however, these data are difficult to standardize given high intra-annual variation in individual mass. Importantly, previous analyses have noted that pup fitness may be only asymptotically related to weaning mass, with survival to reproductive recruitment rising until the observed average mass of 51.5 kg (Bowen et al. 2006), and even slightly decreasing at the highest weaning masses (Bowen et al. 2015). So, assessment of female quality should not rely solely on offspring weaning mass.

After the collapse of the cod fishery in the early 1990s, there was an increase in the pelagic forage fish on the Scotian Shelf (Frank et al. 2005). Since the mid-2000s, silver hake (**?**) and redfish (DFO 2018a) are among grey seal prey that are at high or increasing abundance, while other prey such as herring (DFO 2017) and cod (DFO 2018b) remain at low levels. In the context of our results, high performance in later age-class may be subsidized by this surge of forage during later years. While most individuals decreased in reproductive performance at this time, the increase in prey base may partially explain the magnified variation in performance; high quality individuals may have been able to secure ideal foraging habitats to capitalize on the food increase that may then be relatively unavailable to poorer competitors (van Noordwijk and de Jong 1986, Plard et al. 2014).

With its roots in *r*-vs. *K*-selection (MacArthur and Wilson 1967, Pianka 1970), pace-of-life syndrome has suggested much of life-history variation falls on a slow–fast continuum, with low reproductive rate, slow development and long life span at one end and rapid development, short lifespans, and high reproductive rates at the other end. While this has been well established among a variety of taxa (Wikelski et al. 2003, Wiersma et al. 2007, Hamel et al. 2010), we found no support for POLS among individuals within the increasing Sable Island breeding population of grey seals. Because grey seals are top predators and experience virtually no predation, especially as adults, extrinsic mortality associated with prolonged foraging is likely nearly zero. Providing digestion and energy storage and subsequent mobilization of resources during lactation incurs relatively minimal somatic wear (Zhang and Hood 2016), allowing individuals to invest heavily in each reproductive attempt such that predictions from POLS may not be applicable (Reznick 1982). Of course, the POLS hypothesis is not only concerned with the pace of reproduction, but also a suite of physiological (e.g. metabolic, hormonal, immunological) and behavioral traits that have coevolved with life histories and constrain individual responses to environmental stresses and perceived risks (Ricklefs and Wikelski 2002, Réale et al. 2010).

### Effect of age class reproductive performance on survival

Likely, the positive association of lifetime reproductive investment with survival rates is largely driven by late life investment, as other age classes have more modest effects across model fits (Table 2, Appendix S1: Table S3). In both reproductive frequency and provisioning performance, late life reproductive investment was most strongly correlated with survival (Table 2, Appendix S1: Table S3), and this age class also had the greatest variation among individuals across traits. The predominant hypotheses concerning the evolution of senescence hinge upon the declining force of selection with age (Williams 1957, Kirkwood 1977), such that individual fitness is most sensitive to changes in early reproduction (Forslund and Part 1995, Nussey et al. 2013). Thus, individual reproductive variation could be smaller during early reproduction, when the force of selection is strongest.

Individual quality as a driving force in life history evolution has been demonstrated in previous analyses as positive associations between early reproductive traits (age and success at first reproduction) and future breeding/longevity (Weladji et al. 2008, Bonenfant et al. 2009, Hamel et al. 2009, Aubry et al. 2011, Fay et al. 2016). Many of these studies did not include late-life reproductive traits, and results vary among the few that did. In short-tailed shearwaters, for example, early reproductive performance was positively associated with longer lifespans, but late life performance increased mortality risk (Wooller et al. 1990). Aubry et al. (2011) found that, in black-legged kittiwakes, there is an immediate survival cost to reproduction (attempting to breed at time *t* increased mortality risk to time *t* + 1), but early reproducers and individuals with more breeding experience lived longer. By integrating lifetime and age class specific reproductive performance to explore relationships between reproductive investment and longevity, we find that in our system early reproductive traits do not have notable impacts on future survival or breeding success. Instead, our results point to late life reproductive achievement as an indicator of female quality. We found that late life reproductive performance was strongly correlated with previous performance and was also the primary driver of the positive association between reproductive success and survival. These results could be the result of individual variation in senescence and selective disappearance of poor-quality mothers (such as in Cam et al. 2002, Mauck et al. 2004), so further investigation into age-related reproductive variation among individuals may yield interesting insights into the evolution of reproductive tactics and their impacts on population dynamics (Hamel et al. 2018).

Interestingly, the two reproductive traits differed in how consistently individuals performed relative to other females across age classes. Our models estimated a striking consistency in individual provisioning performance across age classes relative to other females, and substantially weaker correlation among age classes in reproductive frequency (Figure 2). Previous analyses noted that there is also greater variation among individuals in reproductive frequency (Badger et al. 2020), which may be explained by the nature of the data (discrete and more prone to observation error), but also perhaps reproductive frequency is a more plastic trait than provisioning performance. Long-lived, iteroparous species such as grey seals have high survival rates likely because they are able to vary reproductive effort in response to environmental conditions (Bull and Shine 1979). With limited resources, females may be more likely to forego reproduction altogether than support gestation and birth only to provide insufficient or highly variable care to dependent offspring. Due to innate physiological differences among individuals, females may consistently deliver a higher or lower amount of provisions to young than others, retaining their ranking across age classes (Lang et al. 2009). Consequently, although there are age-related changes in provisioning (Bowen et al. 2006), the hierarchy of individuals in provisioning performance is generally maintained as cohorts age. This result supports the conclusion that individual quality drives differences among individuals in reproductive performance of female grey seals.

### The structure of heterogeneity and consequences for population dynamics

Individual variation in reproductive performance and survival probability affect both deterministic and stochastic properties of population processes, driving population plasticity and evolution in response to environmental change (Vindenes et al. 2008, Paterson et al. 2018). Heterogeneity can alter the magnitude of demographic stochasticity thus affecting population dynamics and persistence (Lomnicki 1978, Conner and White 1999, Stover et al. 2012). When vital rates are positively correlated at an individual level, as in our study, demographic stochasticity may have an even larger impact particularly at small population sizes, because individuals are not homogeneous: the loss of a “high quality” female would have more profound effects on population recovery, as she would survive to reproduce more offspring than a “poor quality” female (Conner and White 1999, Badger et al. 2020). Positive associations of survival and reproductive abilities also mean that phenotypic selection within birth cohorts will lead to a decrease in the proportion of individuals with lower survival and correspondingly lower reproductive performance, which can mask senescence (Vaupel and Yashin 1985, Forslund and Part 1995, Cam et al. 2002). Such masking could be exacerbated by increased intraspecific competition, such as a consequence of increased population density. These effects are often not included in population dynamics models or projections, and even when they are, there is often no data to constrain model parameters (Vindenes et al. 2008).

Individual heterogeneity is ubiquitous in natural populations, but the extent to which inter-individual variation structures life history patterns versus energetic trade-offs has become a lively debate in ecology (Bergeron et al. 2011, Cam et al. 2016, Gimenez et al. 2017). Nevertheless, few studies have simultaneously examined these hypotheses, and some of those arbitrarily defined individual quality as single traits such as laying date (reviewed in Moyes et al. 2009). In many other large iteroparous mammals, studies have shown support for trade-offs as a dominating force in life history variation. For example, Moyes et al. (2009) found that, though there are consistent individual differences, a high proportion of variance in longevity and various reproductive measures (high calf birth weight and survival, earlier primiparity, etc.) is not explained by individual quality, and instead trade-offs may explain observed life history tactics in red deer. van Noordwijk and de Jong (1986) suggest that the governing force of life history variation may be a result of the interaction between resource acquisition and allocation. If individuals vary more in their ability to acquire resources than how they allocate those resources to competing functions, individual quality may be more important to the observed distribution of life histories because some individuals can invest more in all functions. Alternatively, if all individuals obtain similar resources, then their strategy to allocate resources (i.e. through trade-offs) may contribute more to observed history variation. Consistent individual differences in fitness traits (i.e. quality) and trade-offs among fitness components simply constitute two sources of variation in the distribution of life histories, and the relative magnitude and interaction of these sources will determine important processes in population dynamics and evolutionary patterns.

## Conclusions & Implications

Our results revealed that individuals in the Sable Island breeding colony that reproduce more often invest more in offspring and survive longer. During much of our study female’s reproductive lives, this population has been experiencing slowing growth as the population presumably is reaching carrying capacity (Bowen 2011, den Heyer et al. 2017). Evidence of the effects of negative density dependence thus far include a drop in juvenile apparent survival to reproductive recruitment thought to stem from increased competition with adults for high quality foraging habitat (den Heyer et al. 2013, Breed et al. 2013). Previous work showed that the increasing density of seals on the Scotian shelf is associated with an increase in differences among individuals in reproductive performance (Badger et al. 2020), we show here that individual reproductive performance is also positively correlated with survival. As population size increases and competition intensifies, we could expect differences in reproductive abilities and survival to become more pronounced, potentially resulting in the proportion of high quality individuals to grow and the population-level vital rates to increase as individuals with lower reproductive capabilities and correspondingly lower survival disappear from the breeding colony.

We note that our sampling scheme and modeling framework may yield only a conservative estimate of the relationship between survival and reproductive performance, as we only included individuals that (1) had been observed on the island on at least 2 occasions and (2) nursed their pup long enough to be recorded by our research teams. These qualifications result in a sample that only explores part of the spectrum of reproductive investment and survivability. Compared to previous robust survival analyses using a larger subset of these data, our survival estimates are biased slightly high for younger females (aged 4-24, 0.993 ± 0.002 here vs. 0.989±0.001 in den Heyer and Bowen 2017). Inexperienced or low quality mothers may frequently flee or abandon pups when disturbed and these reproductive attempts would not be recorded in our observations, though this is not a source of serious bias (Hammill et al. 2017). Pups of skittish females that are found alone may be deemed weaned, and if these females returned to nurse and remained undetected from research teams, her provisioning performance would be underrepresented. Additionally, individuals that do not recruit to the breeding population or die early in life would not be sampled. For these reasons, poor performers are much less likely to be observed, resulting in a larger proportion of high quality females in our sample than present in the Sable Island breeding population. Sex-specific adult grey seal survival has been estimated for the population assessment model (den Heyer et al. 2013, den Heyer and Bowen 2017), so our goal was not to estimate survival here, only to understand the relationship between survival and reproductive allocation.

## Appendix S1

## References

Araújo MS, Bolnick DI, Layman CA. 2011. The ecological causes of individual specialisation. Ecology Letters 14: 948–958. doi: 10.1007/s11661-997-0172-9.

Arnbom T, Fedak MA, Boyd IL. 1997. Factors affecting maternal expenditure in southern elephant seals during lactation. Ecology 78: 471–483. doi: 10.1890/0012-9658(1997)078[0471:FAMEIS]2.0.CO;2.

Aubry LM, Cam E, Koons DN, Monnat JY, Pavard S. 2011. Drivers of age-specific survival in a long-lived seabird: Contributions of observed and hidden sources of heterogeneity. Journal of Animal Ecology 80: 375–383. doi: 10.1111/j.1365-2656.2010.01784.x.

Aubry LM, Koons DN Monnat and JY Emmanuelle C. 2009. Consequences of recruitment decisions and heterogeneity on age-specific breeding success in a long-lived seabird. Ecology 90: 2491–2502. doi: 10.1890/08-1475.1.

Austin D, Bowen WD, McMillan JI. 2004. Intraspecific variation in movement patterns: Modeling individual behaviour in a large marine predator. Oikos 105: 15–30. doi: 10.1111/j.0030-1299.1999.12730.x.

Badger JJ, Bowen WD, den Heyer CE, Breed GA. 2020. Variation in individal reproductive performance amplified with population size in a long-lived carnivore. Ecology e03024. doi: 10.1002/ecy.3024.

Beauplet G, Barbraud C, Dabin W, Kussener C, Guinet C, Benton T. 2006. Age-specific survival and reproductive performances in fur seals: Evidence of senescence and individual quality. Oikos 112: 430–441. doi: 10.1111/j.0030-1299.2006.14412.x.

Begon M, Parker GA. 1986. Should egg size and clutch size decrease with age ? Oikos 47: 293–302. doi: 10.2307/3565440.

Bell G. 1980. The costs of reproduction and their consequences. American Naturalist 116: 45–76. doi: 10.1086/283611.

Bergeron P, Baeta R, Pelletier F, Réale D, Garant D. 2011. Individual quality: Tautology or biological reality? Journal of Animal Ecology 80: 361–364. doi: 10.1111/j.1365-2656.2010.01770.x.

Bérubé CH, Festa-Bianchet M, Jorgenson JT. 1999. Individual differences, longevity, and reproductive senescence in bighorn ewes. Ecology 80: 2555–2565. doi: 10.2307/177240.

Blomberg EJ, Sedinger JS, Nonne DV, Atamian MT. 2013. Seasonal reproductive costs contribute to reduced survival of female greater sage-grouse. Journal of Avian Biology 44: 149–158. doi: 10.1111/j.1600-048X.2012.00013.x.

Blomquist GE. 2009. Trade-off between age of first reproduction and survival in a female primate. Biology Letters 5: 339–342. doi: 10.1098/rsbl.2009.0009.

Bolnick DI, Amarasekare P, Araújo MS, Bürger R, Levine JM, Novak M, Rudolf VHW, Schreiber SJ, Urban MC, Vasseur DA. 2011. Why intraspecific trait variation matters in community ecology. Trends in Ecology and Evolution 26: 183–192. doi: 10.1016/j.tree.2011.01.009.

Bolnick DI, Svanbäck R, Fordyce JA, Yang LH, Davis JM, Hulsey CD, Forister ML. 2003. The ecology of individuals: incidence and implications of individual specialization. American Naturalist 161: 1–28. doi: 10.1086/343878.

Bonenfant C, Pelletier F, Garel M, Bergeron P. 2009. Age-dependent relationship between horn growth and survival in wild sheep. Journal of Animal Ecology 78: 161–171. doi: 10.1111/j.1365-2656.2008.01477.x.

Boness DJ, James H. 1979. Reproductive behaviour of the grey seal (Halichoerus grypus) on Sable Island, Nova Scotia. Journal of Zoology 188: 477–500. doi: 10.1111/j.1469-7998.1979.tb03430.x.

Bowen WD. 2011. Historical grey seal abundance and changes in the abundance of grey seal predators in the Northwest Atlantic. DFO Canadian Science Advisory Secretariat Research Document 2011/026.

Bowen WD, den Heyer CE, McMillan JI, Iverson SJ. 2015. Offspring size at weaning affects survival to recruitment and reproductive performance of primiparous gray seals. Ecology and Evolution 5: 1412–1424. doi: 10.1002/ece3.1450.

Bowen WD, Iverson SJ, McMillan JI, Boness DJ. 2006. Reproductive performance in grey seals: Age-related improvement and senescence in a capital breeder. Journal of Animal Ecology 75: 1340–1351. doi: 10.1111/j.1365-2656.2006.01157.x.

Bowen WD, McMillan JI, Blanchard W. 2007. Reduced population growth of gray seals at Sable Island: Evidence from pup production and age of primiparity. Marine Mammal Science 23: 48–64. doi: 10.1111/j.1748-7692.2006.00085.x.

Bowen WD, Stobo WT, Smith SJ. 1992. Mass changes of grey seal Halichoerus grypus pups on Sable Island: differential maternal investment reconsidered. Journal of Zoology, London 227: 607–622. doi: 10.1111/j.1469-7998.1992.tb04418.x.

Breed GA, Bowen WD, Leonard ML. 2013. Behavioral signature of intraspecific competition and density dependence in colony-breeding marine predators. Ecology and Evolution 3: 3838–3854. doi: 10.1002/ece3.754.

Breed GA, Jonsen ID, Myers RA, Bowen WD, Leonard ML, Bowen WD, Leonard ML. 2009. Sex-specific, seasonal foraging tactics of adult grey seals (Halichoerus grypus) revealed by state-space analysis. Ecology 90: 3209–3221. doi: 10.1890/07-1483.1.

Brooks SP, Gelman A. 1998. General methods for monitoring convergence of iterative simulations. Journal of Computational and Graphical Statistics 7: 434–455. doi: 10.1080/10618600.1998.10474787.

Bull JJ, Shine R. 1979. Iteroparous animals that skip opportunities for reproduction. American Naturalist 114: 296–303.

Cam E, Aubry LM, Authier M. 2016. The conundrum of heterogeneities in life history studies. Trends in Ecology and Evolution 31: 872–886. doi: 10.1016/j.tree.2016.08.002.

Cam E, Link WA, Cooch EG, Monnat JY, Danchin E. 2002. Individual covariation in life-history traits: seeing the trees despite the forest. American Naturalist 159: 96–105. doi: 10.1086/324126.

Chambert T, Rotella JJ, Higgs MD, Garrott RA. 2013. Individual heterogeneity in reproductive rates and cost of reproduction in a long-lived vertebrate. Ecology and Evolution 3: 2047–2060. doi: 10.1002/ece3.615.

Charmantier A, Perrins C, McCleery RH, Sheldon BC. 2006. Quantitative genetics of age at reproduction in wild swans: Support for antagonistic pleiotropy models of senescence. Proceedings of the National Academy of Sciences 103: 6587–6592. doi: 10.1073/pnas.0511123103.

Clutton-Brock TH. 1984. Reproductive effort and terminal investment in iteroparous animals. American Naturalist 123: 212–229. doi: 10.1086/284198.

Clutton-Brock TH, Albon SD, Guinness FE. 1989. Fitness costs of gestation and lactation in wild mammals. Nature 337: 260–262. doi: 10.1098/rsbl.2016.0417.

Cody ML. 1966. A general theory of clutch size. Evolution 20: 174–184. doi: 10.2307/2406571.

Conner MM, White GC. 1999. Effects of individual heterogeneity in estimating the persistence of small populations. Natural Resource Modeling 12: 109–127. doi: 10.1111/j.1939-7445.1999.tb00005.x.

Cox RM, Calsbeek R. 2010. Severe costs of reproduction persist in Anolis lizards despite the evolution of a single-egg clutch. Evolution 64: 1321–1330. doi: 10.1111/j.1558-5646.2009.00906.x.

Crocker DE, Champagne CD, Fowler MA, Houser DS. 2014. Adiposity and fat metabolism in lactating and fasting northern elephant seals. Advances in Nutrition 5: 57–64. doi: 10.3945/an.113.004663.

Curio E. 1983. Why do young birds reproduce less well? Ibis 125: 400–404. doi: 10.1111/j.1474-919X.1983.tb03130.x.

De Loof A. 2011. Longevity and aging in insects: Is reproduction costly; cheap; beneficial or irrelevant? A critical evaluation of the “trade-off” concept. Journal of Insect Physiology 57: 1–11. doi: 10.1016/j.jinsphys.2010.08.018.

den Heyer CE, Bowen WD. 2017. Estimating changes in vital rates of Sable Island grey seals using mark-recapture analysis. Canadian Science Advisory Secretariat Research Document 2017/054.

den Heyer CE, Bowen WD, Mcmillan JI. 2013. Long-term changes in grey seal vital rates at Sable Island estimated from POPAN mark-resighting analysis of branded seals. DFO Canadian Science Advisory Secretariat Research Document 2013/021.

den Heyer CE, Lang SLC, Bowen WD, Hammill MO. 2017. Pup production at Scotian shelf grey seal (Halichoerus grypus) colonies in 2016. DFO Canadian Science Advisory Secretariat Research Document 2017/056.

DFO. 2017. Stock assessment of Canadian Northwest Atlantic grey seals (Halichoerus grypus). Canadian Science Advisory Secretariat Science Advisory Report 2017/045.

DFO. 2018a. Assessment of redfish stocks (Sebastes mentella and S. fasciatus) in units 1 and 2 in 2017. Technical Report June, Canadian Science Advisory Secretariat.

DFO. 2018b. Stock status update of Atlantic cod (Gadus morhua) in NAFO divisions 4X5Yb. Technical Report April, Canadian Science Advisory Secretariat.

Fay R, Barbraud C, Delord K, Weimerskirch H. 2016. Variation in the age of first reproduction: different strategies or individual quality? Ecology 97: 1842–1851. doi: 10.1515/fman-2015-0003.

Festa-Bianchet M, Jorgenson JT. 1998. Selfish mothers: reproductive expenditure and resource availability in bighorn ewes. Behavioral Ecology 9: 144–150. doi: 10.1093/beheco/9.2.144.

Fisher RA. 1930. The genetical theory of natural selection. Oxford: Clarendon Press.

Forslund P, Part T. 1995. Age and reproduction in birds - hypotheses and tests. Trends in Ecology & Evolution 10: 374–378. doi: 10.1016/S0169-5347(00)89141-7.

Frank KT, Petrie B, Choi JS, Leggett WC. 2005. Trophic cascades in a formerly cod-dominated ecosystem. Science 308: 1621–1623. doi: 10.1126/science.1108228.

Gaillard JM, Pontier D, Allainé D, Lebreton JD, Trouvilliez J, Clobert J. 1989. An analysis of demographic tactics in birds and mammals. Oikos 56: 59–76. doi: 10.2307/3566088.

George JC, Bada J, Zeh J, Scott L, Brown SE, O’Hara T, Suydam R. 1999. Age and growth estimates of bowhead whales (Balaena mysticetus) via aspartic acid racemization. Canadian Journal of Zoology 77: 571–580. doi: 10.1139/z99-015.

George JC, Thewissen JGM, Von Duyke A, Breed GA, Suydam R, Sformo T, Person B, Brower H. 2020. Chapter 7: Life History, Growth and Form. In Bowhead Whales: Biology and Conservation. Amsterdam: Elsevier.

Gimenez O, Cam E, Gaillard JM. 2017. Individual heterogeneity and capture–recapture models: what, why and how? Oikos 127: 664–686. doi: 10.1111/oik.04532.

Gimenez O, Rossi V, Choquet R, Dehais C, Doris B, Varella H, Vila JP, Pradel R. 2007. State-space modelling of data on marked individuals. Ecological Modelling 206: 431–438. doi: 10.1016/j.ecolmodel.2007.03.040.

Hamel S, Gaillard JM, Douhard M, Festa-Bianchet M, Pelletier F, Yoccoz NG. 2018. Quantifying individual heterogeneity and its influence on life-history trajectories: different methods for different questions and contexts. Oikos 127: 687–704. doi: 10.1111/oik.04725.

Hamel S, Gaillard JM, Festa-Bianchet M, Côté SD. 2009. Individual quality, early-life conditions, and reproductive success in contrasted populations of large herbivores. Ecology 90: 1981–1995. doi: 10.1890/08-0596.1.

Hamel S, Gaillard JM, Yoccoz NG, Loison A, Bonenfant C, Descamps S. 2010. Fitness costs of reproduction depend on life speed: Empirical evidence from mammalian populations. Ecology Letters 13: 915–935. doi: 10.1111/j.1461-0248.2010.01478.x.

Hamilton WD. 1966. The moulding of senescence by natural selection. Journal of Theoretical Biology 12: 12–45. doi: 10.1016/0022-5193(66)90184-6.

Hammill MO, den Heyer CE, Bowen WD, Lang SLC. 2017. Grey seal population trends in Canadian waters, 1960-2016 and harvest advice. DFO Canadian Science Advisory Secretariat Research Document 2017/052: 1–35.

Hunt BP. 1951. Reproduction of the burrowing mayfly, Hexagenia limbata (Serville) in Michigan. The Florida Entomologist 34: 59–70. doi: 10.2307/3492083.

Iverson SJ, Bowen WD, Boness DJ, Oftedall OT. 1993. Effect maternal size and milk energy output on pup growth in grey seals (Halichoerus grypus). Physiological Zoology 66: 61–88. doi: 10.2307/30158287.

Jones OR, Gaillard JM, Tuljapurkar S, Alho JS, Armitage KB, Becker PH, Bize P, Brommer J, Charmantier A, Charpentier M, Clutton-Brock T, Dobson FS, Festa-Bianchet M, Gustafsson L, Jensen H, Jones CG, Lillandt BG, McCleery R, Merila J, Neuhaus P, Nicoll MAC, Norris K, Oli MK, Pemberton J, Pietiainen H, Ringsby TH, Roulin A, Sæther BE, Setchell JM, Sheldon BC, Thompson PM, Weimerskirch H, Jean Wickings E, Coulson T. 2008. Senescence rates are determined by ranking on the fast-slow life-history continuum. Ecology Letters 11: 1–10. doi: 10.1111/j.1461-0248.2008.01187.x.

Kery M, Schaub M. 2012. Two Further Useful Multistate Capture – Recapture Models : 487–496doi: 10.1016/B978-0-12-387020-9.00016-X.

Kirkwood TBL. 1977. Evolution of ageing. Nature 270: 301–304. doi: 10.1038/270301a0.

Kirkwood TBL, Austad SN. 2000. Why do we age? Nature 408: 233–238. doi: 10.1038/35041682.

Lack D. 1947. The significance of clutch size. Ibis : 302–352 doi: 10.1111/j.1474-919X.1947.tb04155.x.

Lang SLC, Iverson SJ, Bowen WD. 2006. Individual variation in milk composition over lactation in harbour seals (Phoca vitulina) and the potential consequences of intermittent attendance. Canadian Journal of Zoology 83: 1525–1531. doi: 10.1139/z05-149.

Lang SLC, Iverson SJ, Bowen WD. 2009. Repeatability in lactation performance and the consequences for maternal reproductive success in gray seals. Ecology 90: 2513–2523. doi: 10.1890/08-1386.1.

Lang SLC, Iverson SJ, Bowen WD. 2011. The influence of reproductive experience on milk energy output and lactation performance in the grey seal (Halichoerus grypus). PLoS ONE 6: 1–11. doi: 10.1371/journal.pone.0019487.

Lemaître JF, Berger V, Bonenfant C, Douhard M, Gamelon M, Plard F, Gaillard JM. 2015. Early-late life trade-offs and the evolution of ageing in the wild. Proceedings of the Royal Society B 282. doi: 10.1098/rspb.2015.0209.

Lidgard DC, Bowen WD, Iverson SJ. 2020. Sex-differences in fine-scale home-range use in an upper-trophic level marine predator. Movement Ecology 8: 1–16. doi: 10.1186/s40462-020-0196-y.

Lomnicki A. 1978. Individual differences between animals and the natural regulation of their numbers. Journal of Animal Ecology 47: 461–475. doi: 10.2307/3794.

MacArthur RH, Wilson EO. 1967. The theory of island biogeography. Woodstock, Oxfordshire: Princeton University Press.

Macdonald KR, Rotella JJ, Garrott RA, Link WA. 2020. Sources of variation in maternal allocation in a long-lived mammal. Journal of Animal Ecology doi: 10.1111/1365-2656.13243.

Mauck RA, Huntington CE, Grubb TC. 2004. Age-specific reproductive success: evidence for the selection hypothesis. Evolution 58: 880–885. doi: 10.1111/j.0014-3820.2004.tb00419.x.

Mellish JE, Iverson SJ, Bowen WD. 1999. Variation in milk production and lactation performance in grey seals and consequences for pup growth and weaning characteristics. Physiological and Biochemical Zoology 72: 677–690. doi: 10.1086/316708.

Moyes K, Morgan BJT, Morris A, Morris SJ, Clutton-Brock TH, Coulson T. 2009. Exploring individual quality in a wild population of red deer. Journal of Animal Ecology 78: 406–413. doi: 10.1111/j.1365-2656.2008.01497.x.

Nussey DH, Froy H, Lemaitre JF, Gaillard JM, Austad SN. 2013. Senescence in natural populations of animals: Widespread evidence and its implications for bio-gerontology. Ageing Research Reviews 12: 214–225. doi: 10.1016/j.arr.2012.07.004.

Partridge L, Harvey PH. 1988. The ecological context of life history evolution. Science 241: 1449–1455. doi: 10.1126/science.241.4872.1449.

Paterson JT, Rotella JJ, Link WA, Garrott R. 2018. Variation in the vital rates of an Antarctic marine predator: the role of individual heterogeneity. Ecology 99: 2385–2396. doi: 10.1002/ecy.2481.

Pianka ER. 1970. On r-and K-Selection. American Naturalist 104: 592–597. doi: 10.1086/282697.

Plard F, Gaillard JM, Coulson T, Hewison AJM, Delorme D, Warnant C, Nilsen EB, Bonenfant C. 2014. Long-lived and heavier females give birth earlier in roe deer. Ecography 37: 241–249. doi: 10.1111/j.1600-0587.2013.00414.x.

Plummer M. 2003. JAGS: A program for analysis of Bayesian graphical models using Gibbs sampling.

Plummer M. 2018. rjags: Bayesian graphical models using MCMC. R package version 4-8.

R Core Team. 2020. R: A language and environment for statistical computing. URL https://www.r-project.org/

Réale D, Garant D, Humphries MM, Bergeron P, Careau V, Montiglio PO. 2010. Personality and the emergence of the pace-of-life syndrome concept at the population level. Philosophical Transactions of the Royal Society B 365: 4051–4063. doi: 10.1098/rstb.2010.0208.

Reed TE, Kruuk LEB, Wanless S, Frederiksen M, Cunningham EJA, Harris MP. 2008. Reproductive senescence in a long-lived seabird: rates of decline in late-life performance are associated with varying costs of early reproduction. American Naturalist 171: E89–E101. doi: 10.1086/524957.

Reed TE, Robin SW, Schindler DE, Hard JJ, Kinnison MT. 2010. Phenotypic plasticity and population viability: The importance of environmental predictability. Proceedings of the Royal Society B 277: 3391–3400. doi: 10.1098/rspb.2010.0771.

Reznick D. 1982. The impact of predation on life history evolution in Trinidadian guppies: genetic basis of observed life history patterns. Evolution 36: 1236. doi: 10.2307/2408156.

Reznick D, Nunney L, Tessier A. 2000. Big houses, big cars, superfleas and the costs of reproduction. Trends in Ecology and Evolution 15: 421–425. doi: 10.1016/S0169-5347(00)01941-8.

Ricklefs R, Wikelski M. 2002. The physiology/life history nexus. Trends in Ecology & Evolution 17: 462–468.

Roff DA. 1992. The evolution of life histories. New York: Chapman and Hall.

Roff DA. 2002. Life history evolution. Sunderland: Sinaeur Associates.

Royle JA. 2008. Modeling individual effects in the Cormack-Jolly-Seber model: A state-space formulation. Biometrics 64: 364–370. doi: 10.1111/j.1541-0420.2007.00891.x.

Salguero-Gómez R, Jones OR, Jongejans E, Blomberg SP, Hodgson DJ, Mbeau-Ache C, Zuidema PA, De Kroon H, Buckley YM. 2016. Fast-slow continuum and reproductive strategies structure plant life-history variation worldwide. Proceedings of the National Academy of Sciences 113: 230–235. doi: 10.1073/pnas.1506215112.

Solymos P. 2010. dclone: Data cloning in R. The R Journal 2: 29–37.

Stearns SC. 1989. Trade-offs in life history evolution. Functional Ecology 3: 259–268. doi: 10.2307/2389364.

Stearns SC. 1992. The evolution of life histories. Oxford: Oxford University Press.

Stearns SC. 2000. Life history evolution: Successes, limitations, and prospects. Naturwissenschaften 87: 476–486. doi: 10.1007/s001140050763.

Stoelting RE, Gutiérrez RJ, Kendall WL, Peery MZ. 2015. Life-history tradeoffs and reproductive cycles in Spotted Owls. The Auk 132: 46–64. doi: 10.1642/auk-14-98.1.

Stover JP, Kendall BE, Fox GA. 2012. Demographic heterogeneity impacts density-dependent population dynamics. Theoretical Ecology 5: 297–309. doi: 10.1007/s12080-011-0129-x.

Thompson D, Fedak MA. 1993. Cardiac responses of grey seals during diving at sea. Journal of Experimental Biology 174: 139–164.

van Noordwijk AJ, de Jong G. 1986. Acquisition and allocation of resources: their influence on variation in life history tactics. American Naturalist 128: 137–142. doi: 10.1086/284547.

Vaupel JW, Yashin AI. 1985. Heterogeneity’s ruses: some surprising effects of selection on population dynamics. The American Statistician 39: 176–185. doi: 10.1080/00031305.1985.10479424.

Vindenes Y, Engen S, Sæther BE. 2008. Individual heterogeneity in vital parameters and demographic stochasticity. American Naturalist 171: 455–467. doi: 10.1086/528965.

Watanabe S. 2010. Asymptotic equivalence of Bayes cross validation and Widely Applicable Information Criterion in singular learning theory. Journal of Machine Learning Research 11: 3571–3594.

Weitzman J, den Heyer C, Bowen DW. 2017. Factors influencing and consequences of breeding dispersal and habitat choice in female grey seals (Halichoerus grypus) on Sable Island, Nova Scotia. Oecologia 183: 367–378. doi: 10.1007/s00442-016-3764-5.

Weladji RB, Loison A, Gaillard JM, Holand O, Mysterud A, Yoccoz NG, Nieminen M, Stenseth NC. 2008. Heterogeneity in individual quality overrides costs of reproduction in female reindeer. Oecologia 156: 237–247. doi: 10.1007/s00442-008-0961-x.

Wiersma P, Muñoz-Garcia A, Walker A, Williams JB. 2007. Tropical birds have a slow pace of life. Proceedings of the National Academy of Sciences of the United States of America 104: 9340–9345. doi: 10.1073/pnas.0702212104.

Wikelski M, Spinney L, Schelsky W, Scheuerlein A, Gwinner E. 2003. Slow pace of life in tropical sedentary birds: A common-garden experiment on four stonechat populations from different latitudes. Proceedings of the Royal Society B 270: 2383–2388. doi: 10.1098/rspb.2003.2500.

Williams GC. 1957. Pleiotropy, natural selection, and the evolution of senescence. Evolution 11: 398–411. doi: 10.2307/2406060.

Williams GC. 1966. Natural selection, the costs of reproduction, and a refinement of Lack’s principle. American Naturalist 100: 687–690. doi: 10.1086/282461.

Williams TM, Davis RW, Fuiman LA, Francis J, LeBoeuf BJ, Horning M, Calambokidis J, Croll DA. 2000. Sink or swim: strategies for cost-efficient diving by marine mammals. Science 288: 133–136. doi: 10.1126/science.288.5463.133.

Wilson AJ, Nussey DH. 2009. What is individual quality ? An evolutionary perspective. Trends in Ecology & Evolution 25: 207–214. doi: 10.1016/j.tree.2009.10.002.

Wooller RD, Bradley JS, Skira IJ, Serventy DL. 1990. Reproductive success of short-tailed shearwaters Piuffinus tenuirostris in relation to their age and breeding experience. Journal of Animal Ecology 59: 487–496. doi: 10.2307/5165.

Zhang Y, Hood WR. 2016. Current versus future reproduction and longevity: a re-evaluation of predictions and mechanisms. Journal of Experimental Biology 219: 3177–3189. doi: 10.1242/jeb.132183.

